# Telomeric G-Quadruplexes Formed by Six G-overhang Permutation Repeats: Stability, Topology and Potential Physiological Role

**DOI:** 10.64898/2026.07.14.738422

**Authors:** Ziyue Guan, Tengshuo Luo, Jiqing Shen, Shilong Zhang, Chengjie Chu, Jun Zhou, Jean-Louis Mergny, Mingpan Cheng

## Abstract

The single-stranded G-overhangs of vertebrate telomeres, composed of guanine (G)-rich hexanucleotide repeats, can adopt six register-dependent permutations: GGTTAG, GTTAGG, TTAGGG, TAGGGT, AGGGTA, and GGGTTA. Telomeric G-quadruplexes (G4s) formed by the first two repeat permutations, which are the most prevalent at human chromosome ends, have been largely overlooked. Here, G4 formation across all six permutations, each containing 4 to 9 repeats, are systematically investigated under varying cation conditions and molecular crowding environments. Our findings reveal that distinct repeat permutations yield markedly different G4 stabilities and folding topologies, which in turn modulate DNA polymerase activity. This expanded perspective on telomeric structural diversity provides new insights into telomere biology and suggests that multiple repeat patterns may cooperatively influence chromosome end protection and targeting.

## Introduction

Telomeres are specialized nucleoprotein complexes that cap the termini of eukaryotic chromosomes, which are composed of tandem guanine rich DNA repeats bound by the shelterin complex [1,2]. This architecture prevents chromosome ends from being misrecognized as sites of DNA double strand breaks, thereby averting unchecked DNA damage signaling, nucleolytic degradation, and end to end chromosome fusion [3,4]. In vertebrates, as well as a few other species such as some invertebrates and fungi (*e*.*g*., Neurospora crassa), the very end of telomeric DNA consists of hundreds to thousands of 5′-TTAGGG-3′ repeats, terminating in a single stranded G-rich 3′ overhang [5-8]. This G-overhang interacts with telomere binding proteins to modulate telomere length homeostasis and end protection [9,10]. In human somatic cells, progressive telomere attrition during replication is tightly linked to replicative senescence and organismal aging [11,12], whereas telomere maintenance by telomerase or alternative lengthening of telomeres (ALT) pathways is a hallmark of immortalized and malignant cells [13-15].

A distinctive structural feature of G-overhangs is their ability to fold into four stranded G-quadruplex (G4) structures, stabilized by Hoogsteen hydrogen bonding between guanines and monovalent cations such as K^+^ and Na^+^ [16-18]. G4 folding topology, which can be parallel, antiparallel or hybrid, critically influences thermodynamic stability, ligand recognition, protein interaction and regulatory function at chromosome ends [19-22].

For decades, structural and biologically functional studies of human telomeric G4 DNAs have been dominated by work on the 5′-TTAGGG-3′ repeat and closely related flanking base permutations [23,24]. This repeat exhibits remarkable structural polymorphism, folds into distinct G4 topologies depending on the ionic and molecular crowding conditions. In the presence of Na^+^, the canonical 22-mer sequence (*e*.*g*., 143D, 5′-AGGG[TTAGGG]^3^-3′) characteristically folds into a basket-type antiparallel G4 with lateral and diagonal loops [20]. A structural equilibrium undergoes a dramatic shift upon exchanging Na^+^ for physiological K^+^. In K^+^ solutions, the repeats predominantly adopt hybrid conformations, featuring three stacked G-tetrads connected by mixed strand orientations and propeller/lateral loops, as captured in the G4s of 2GKU (5′-TTGGG[TTAGGG]^3^A-3′) and 2JSM (5′-TAGGG[TTAGGG]^3^-3′) [25,26]. Furthermore, under dehydrating or molecular crowding conditions that mimic the intracellular environment (*e*.*g*., K^+^ supplemented with PEG-200), the structural ensemble is driven toward a parallel topology with propeller loops, exemplified by the structures of 1KF1 (5′-AGGG[TTAGGG]^3^-3′) and 2LD8 (5′-TAGGG[TTAGGG]^3^-3′) [27,28]. This topological plasticity, shifting from antiparallel (Na^+^) to hybrid (K^+^) and to parallel (K^+^ with crowding agent) in highly homologous 5′-TTAGGG-3′ repeats, provides a critical foundation for understanding dynamic interconversion of telomeric G4s in vivo. Table S1 listed the currently resolved human telomeric G4 structures.

Beyond their inherent structural polymorphism, the folding of telomeric overhangs into G4s is intricately linked to their biological functions, particularly in mediating critical protein-DNA interactions at the chromosome ends [29-31]. A primary example is the maintenance of telomere length by telomerase, a specialized ribonucleoprotein that adds the hexanucleotide repeats to the 3’ terminus of telomere [32]. The spontaneous folding of single-stranded G-overhang into stable G4s generally acts as a steric barrier, which shields the terminal 5′-TTAGGG-3′ substrate from being recognized, thereby effectively inhibiting telomerase-mediated elongation [33-36]. Consequently, to facilitate continuous telomere extension and unimpeded DNA replication, specialized DNA helicases or proteins, such as POT1, TPP1, RTEL1, BLM, WRN and others are required to resolve the G4 structures and restore an accessible linear state [37-39].

Given this endogenous regulatory mechanism, telomeric G4s have emerged as compelling therapeutic targets [40]. Over the past decades, numerous small-molecule ligands have been developed to selectively bind and stabilize the canonical G4s formed by 5′-TTAGGG-3′ repeats. These ligands were initially designed as indirect telomerase inhibitors and shown to interfere with telomeric functions, triggering acute telomere uncapping and apoptosis in malignant cells [41-47]. However, these compounds were also active on telomerase-negative tumors: approximately 10-15% of human tumors maintain telomere length through a telomerase-independent recombination mechanism known as ALT [48,49]. Evidence indicates that the stabilization of telomeric G4s and R-loops synergistically upregulates ALT activity [50,51]. This identifies G4 structures as novel targets for ALT-positive malignancies. Consequently, G4 structures have emerged as specific therapeutic vulnerabilities for ALT-positive malignancies, and several telomeric G4 stabilizers show greater activity on telomerase-negative tumors [52,53].

Recent telomere length measurements with nucleotide resolution reveal that 5′-GGTTAG-3′ (61%) and 5′-GTTAGG-3′ (13%) are the most abundant sequences at the G-overhang terminus, significantly exceeding the frequencies of other permutations, including 5′-TTAGGG-3′ (4%), 5′-TAGGGT-3′ (7%), 5′-AGGGTT-3′ (7%), and 5′-GGGTTA-3′ (7%) (Figure 1) [54]. Telomerase employs its 11-nucleotide RNA template (5′-CUAACCCUAAC-3′) to achieve repeat addition processivity, enabling the iterative synthesis of the telomeric repeat 5′-GGTTAG-3′ onto a DNA substrate, explaining why this motif is by far the most frequent at the terminus [55,56]. Notably, all six aforementioned G-overhang terminal permutations can serve as substrates for telomerase, albeit with distinct binding affinities and catalytic efficiencies [57,58]. Surprisingly, although human telomeric G4s are the most extensively studied G4 structures, research on their formation and function has largely focused on the least prevalent G-overhang terminus permutations (Figure 1 and Table S1) [59-62]. This bias likely stems from the fact that these four minor repeat permutations contain all contiguous G-tracts (GGG), which may facilitate a classical G4 folding. In contrast, the two most abundant repeat permutations harbor a split G-tract, a structural feature that may explain why they have been long overlooked.

**Figure 1.**
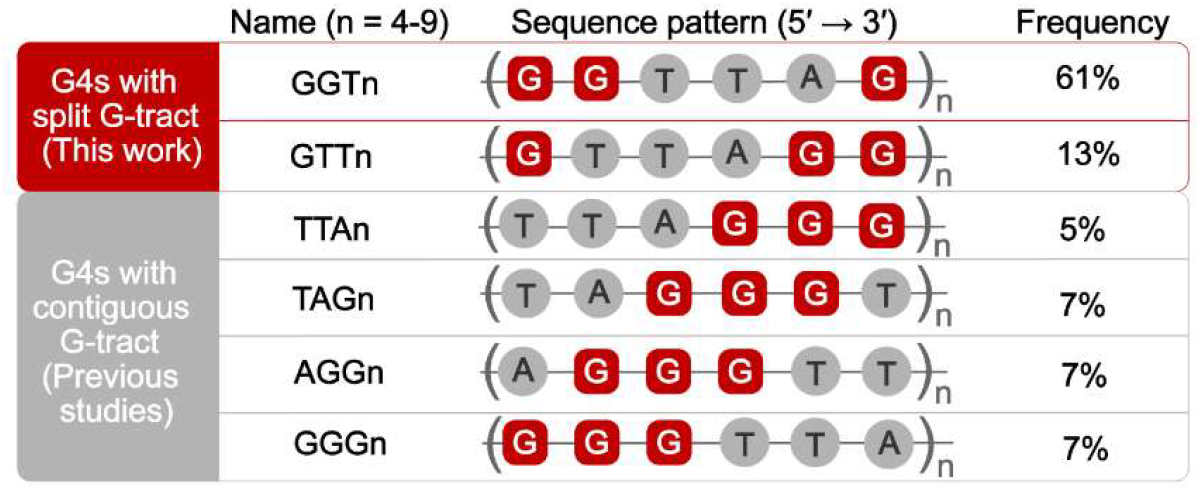
G4s formation by the six telomeric G-overhang permutation repeats: nomenclature, sequences and distribution percentage in 3′ terminus of telomere.

Despite their physiological dominance, the structural and functional consequences of these split G-tract permutations remain largely unresolved, creating a critical blind spot in our comprehension of chromosome end biology. To resolve the current research bias and reveal the authentic properties of human telomeres, G4 formation by all six permutations of the G-overhang terminus, each containing 4 to 9 repeats, was analyzed in this study in the presence of K^+^ and/or Na^+^, without and with a molecular crowding agent (Figure 1). Our findings highlight that subtle variations in the base order of the hexameric repeat units can lead to marked changes in G4 stability, folding topology and DNA polymerase activity. By comprehensively characterizing these previously overlooked sequences, this work establishes an expanded perspective on telomeric structural diversity, with direct implications for telomere biology, regulation, and potential therapeutic targeting.

## Results

### Assessment of telomeric G4s formation

The six G-overhang permutations, named by their first three nucleotide (*e*.*g*., GGTn for (GGTTAG)_n_), have repeat numbers n from 4 to 9, as defined in Figure 1. The total number of sequences is 36 (sequences given in Table S2). The following experiments were performed under eight buffers, designated as follows: 140 mM K^+^ (K^+^), 140 mM Na^+^ (Na^+^), 100 mM K^+^ + 40 mM Na^+^ (K^+^+Na^+^), and 100 mM Na^+^ + 40 mM K^+^ (Na^+^+K^+^), each tested in the absence and presence of 40% (w/v) PEG-200 to mimic a molecular crowding environment. Thermal difference spectra (TDS) indicate that all 36 sequences form G4 structures across all eight buffer conditions (Figure 2A, Figures S1-S6). For example, the TDS profile of the four-repeat G-overhang permutation in K^+^ buffer shows characteristic G4 signals: positive peaks between 245-270 nm and a pronounced negative peak at ∼295 nm [63] (Figure 2A).

**Figure 2.**
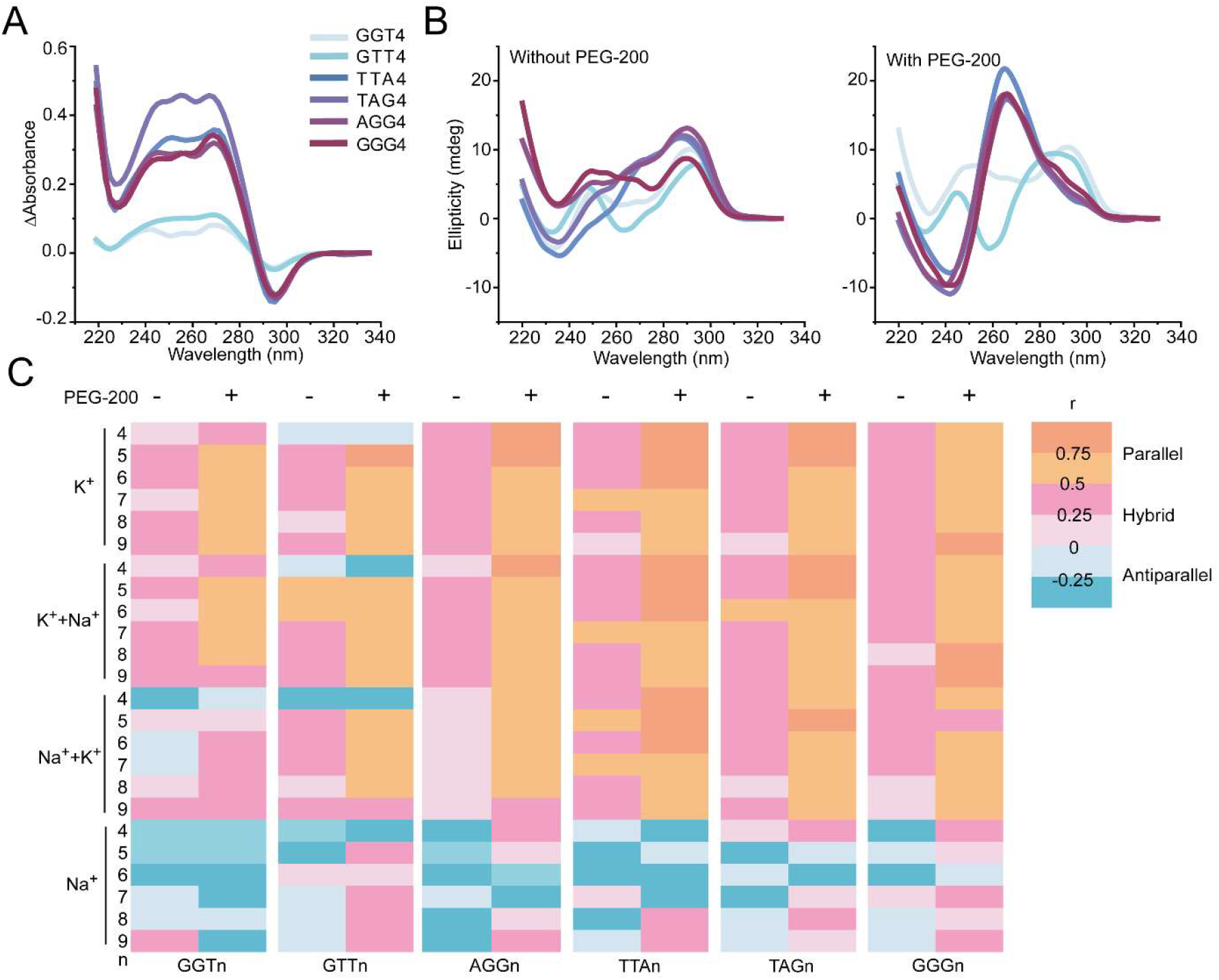
Folding topology of telomeric G4s. (A) TDS of GGT4, GTT4, TTA4, TAG4, AGG4, and GGG4 in 140 mM K^+^. (B) CD spectra in 140 mM K^+^ without (left) and with (right) 40% (w/v) PEG-200 at 25 °C. (C) Heatmap of r values derived from CD spectra measured in 140 mM K^+^ (K^+^), 100 mM K^+^ + 40 mM Na^+^ (K^+^ + Na^+^), 100 mM Na^+^ + 40 mM K^+^ (Na^+^ + K^+^), and 140 mM Na^+^ (Na^+^), without (-) or with (+) PEG-200. r values were calculated as described in Methods; color corresponds to topology assignment: r < 0, antiparallel; 0 ≤ r ≤ 0.5, hybrid; r > 0.5, parallel.

Fluorescence probe binding assays with thioflavin T (ThT) and N-methyl mesoporphyrin IX (NMM) further support the above finding [64]. In K^+^-containing buffers, all telomeric repeats induced strong fluorescence enhancements for both probes, confirming G4 formation (Figures S7 and S8). In contrast, fluorescence enhancements in Na^+^ buffer were limited, particularly for NMM, likely due to the distinct binding selectivity of ThT and NMM for different G4 topologies [65,66]. Conversely, in the presence of 40% PEG-200, the DNA-induced fluorescence enhancement of both probes was severely suppressed (data not shown).

### Folding topologies of telomeric G4s

CD spectroscopy was applied to check the G4 topologies [67]. In K^+^-containing buffer, most G4s exhibited a characteristic spectrum for the hybrid conformation, with a negative peak near 235 nm and a positive peak around 290 nm (Figure 2B and Figures S9-16). When 40% PEG-200 was added to introduce crowding, spectra of most sequences shifted toward the classic parallel G4 signature, showing a negative peak at ∼241 nm and a strong positive peak around 265 nm. Notably, GTT4 displayed a distinct behavior in both conditions: in the absence of crowding, it showed a positive band at ∼245 nm, a negative band near 265 nm, and a secondary positive band close to 290 nm, which consistent with an antiparallel topology. Moreover, even under crowding conditions across all ionic environments, GTT4 retained this antiparallel topology. In addition, G4 formed by TTA7 adopted a parallel topology whereas other TTAn (n = 4, 5, 6, 8, 9) predominantly formed hybrid G4s in K^+^-containing buffer. The topology classification heatmap derived from r values calculated from the CD spectra (see Materials and methods) depicts the folding preferences across ionic environments (Figure 2C). In either K^+^ or Na^+^ buffer, addition of PEG-200 increases the value of r parameters, suggesting that it promote a transition from hybrid to parallel or from antiparallel to hybrid topologies (Figure S17). Nevertheless, a few exceptions occurred, primarily when Na^+^ were predominant, in these cases, the decreased r values indicate that PEG-200 unexpectedly further stabilized the antiparallel topologies.

### Thermal stability and stepwise transitions of telomeric G4s

Then the G4 thermal stability for all 36 sequences were measured under eight buffers. UV-melting and annealing curves were provided Figure 3A and Figures S18-S23 and melting temperature (T_m_) are summarized in Figure 2B and Table S3. Although both GGTn and GTTn contain a split G-tract, only the T_m_ of the latter one is significantly lower than that of the other five G-overhang permutations (Figure 3B). For example, in 140 mM K^+^, the T_m_ value of GTT4 is 49.3 °C, while the T_m_ values of GGT4, TTA4, TAG4, AGG4, and GGG4 are 63.6, 61.0, 65.0, 65.1, and 67.9 °C, respectively. Interestingly, GGT4 is more stable than expected.

**Figure 3.**
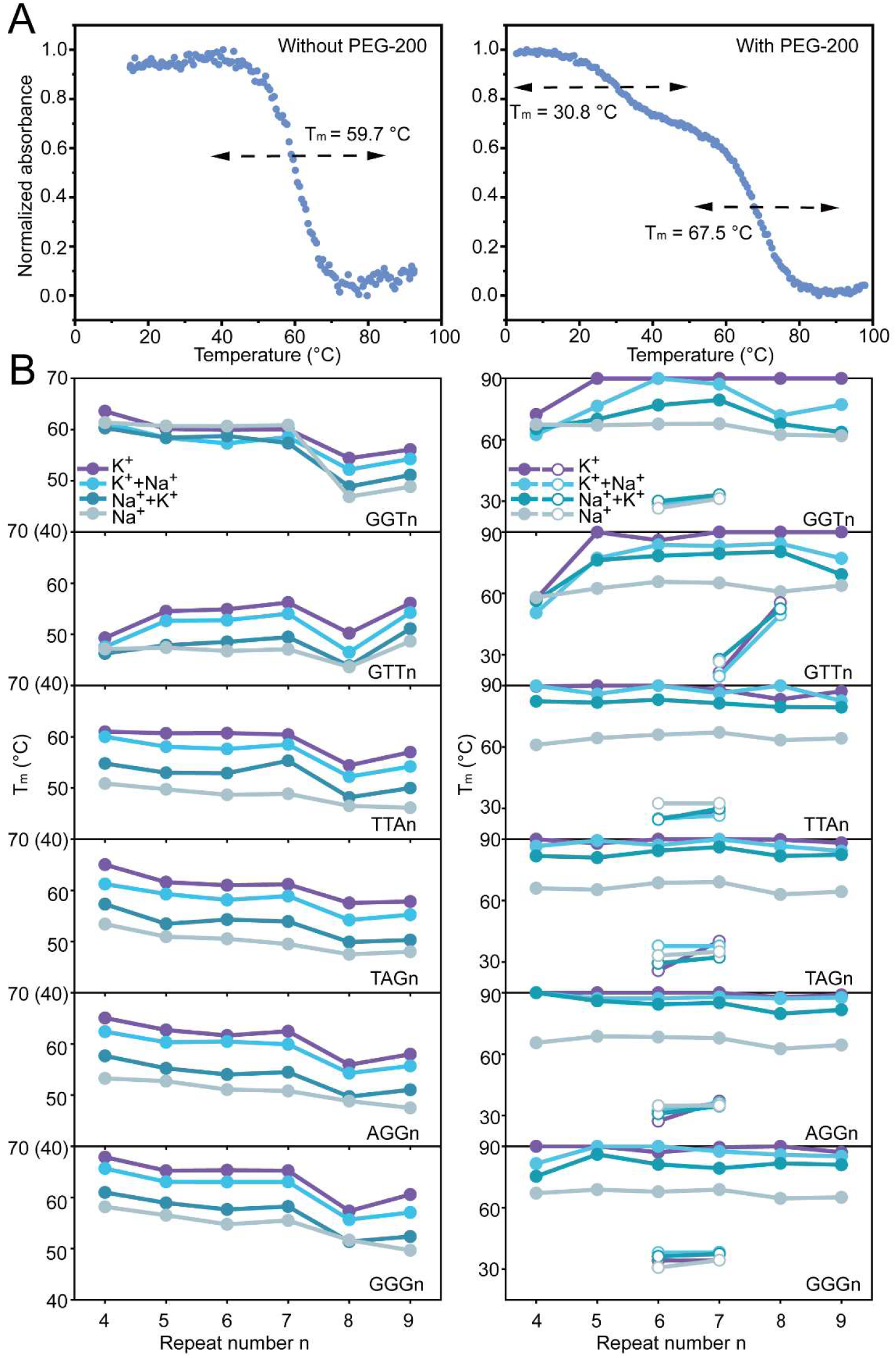
Thermal stability of telomeric G4s. (A) Melting curves of the GGT7 in 140 mM K^+^. (B) Melting temperatures (T_m_) of G-overhang permutation repeats without (left) and with (right) PEG-200. Solid circles represent the higher T_m_ values (right and left), whereas open circles indicate the lower T_m_ values (right). The T_m_ exceeding 90 °C cannot be measured and they are therefore shown as 90 °C here.

Generally, most G-overhang end permutations showed a trend of gradually decreasing T_m_ as the number of tandem repeats increased, indicating that longer overhangs tend to reduce G4 stability (Figure 3B), as previously reported [68]. In addition, also in line with previous reports [67-70], addition of crowding agents greatly enhanced G4 stability. For most groups of permutation repeats, T_m_ values exhibited a local minimum at n = 8, particularly pronounced for GTTn. The recurrence of these inflection points under multiple ionic environments suggests the possibility that specific structural rearrangements may be selectively favored at these repeat numbers. When increasing the repeat number from 4 to 5, the T_m_ values of G4s formed by five of the six permutations (GGTn, TTAn, TAGn, AGGn and GGGn) decreased in the absence of PEG-200, which is consistent with our previous report showing that extra flanking sequences decrease G4 stability [71] (Figure 3B, left). Conversely, the T_m_ value for GTT5 is higher than that for GTT4 (*e*.*g*., 54.5 °C vs. 49.3 °C in K^+^ buffer), suggesting that more guanines participate in G-quartet formation in the former. Similar stability enhancements are observed when the repeat number shifts from n = 4 to n = 5 for both GTTn and GGTn under crowding conditions (Figure 3B, right).

For some G-overhang permutations, two stepwise transitions in both melting and annealing processes appeared in the presence of PEG-200. As a representative example, the UV-melting profiles of GGT7 are shown here to illustrate this unfolding behavior. In the absence of PEG-200, GGT7 displayed a monophasic melting curve characterized by a single unfolding transition with a T_m_ of 59.7 °C (Figure 3A, left), consistent with the denaturation of one predominant G4 structure. Upon addition of PEG-200, the melting curve became clearly biphasic, revealing two stepwise transitions with two T_m_ values of 30.8 and 67.5 °C, respectively (Figure 3A, right). The higher temperature transition may correspond to the same structural element that melted at 59.7 °C without crowding. The lower temperature transition may represent a newly emergent melting event, likely reflecting the unfolding of an alternative folding structure induced specifically under molecular crowding. This additional transition suggests that PEG-200 not only reinforces the stability of preexisting G4s but may also promote the formation of distinct structures, thereby increasing the conformational heterogeneity (Figures S18-23). Interestingly, the PEG-200 induced biphasic melting behavior was limited to a range of repeat numbers for most G-overhang types: for five of the six permutations (GGTn, TTAn, TAGn, AGGn and GGGn), it was only observed at n = 6 and 7; the GTTn displayed a biphasic melting at n = 7 and 8 (Figure 3B, right). Moreover, the T_m_ values of the low-temperature transition differed markedly between GTT7 and GTT8 (ΔT_m_ ≈ 30 °C), whereas in the other five permutations, changes in the T_m_ values of the low-temperature transition between n = 6 and n = 7 were generally small.

### Molecularity of telomeric G4s

Size-exclusion high performance liquid chromatography (SE-HPLC, Figures S24-26) and non-denaturing PAGE (Figures S27-30) were used to reveal the monomeric and multimeric aggregation states of telomeric G4s under defined conditions. GGTn permutation was taken as an example. SE-HPLC showed GGT4 eluted as a single peak at a volume consistent with a monomeric species in both K^+^ and Na^+^ buffers at a concentration of 30 µM (Figure 4A). In contrast, GGT7 showed an additional, earlier eluting peak consistent with a higher order assembly, likely a dimer. This dimer peak was more prominent in K^+^ than in Na^+^, suggesting that dimer formation in GGT7 is favored by potassium. Furthermore, quantitative peak area analysis (Figure S26) revealed a marked increase in the multimeric fraction of GGT7 from 4.2% at 3 µM to 21.5% at 30 µM, in agreement with a concentration-dependent higher-order assembly. Non-denaturing PAGE supported these results (Figure 4B, Figure S27-30). The proportion of monomer and multimer species for all 36 sequences under varying ionic and crowding conditions are summarized in Figure 3C, based on band density analysis of non-denaturing gels. Overall, G4s formed by G-overhang permutations with 4, 5, 8 and 9-repeats folded exclusively into monomeric structure while 7-repeat sequences adopted major monomeric and minor multimeric structures (Figure 4C). Notably, a faint, slower-migrating minor band was observed for GGT8 in both K^+^ and Na^+^ buffers (Figure 4B), however, its close proximity to the primary band suggests that it represents an alternative monomeric conformational species rather than a multimeric assembly. For the G-overhang permutations with 6 repeats, PEG-200 induced the formation of multimeric structures for GGT6, TTA6, TAG6 and GGG6 under some ionic conditions (Figure 4C).

**Figure 4.**
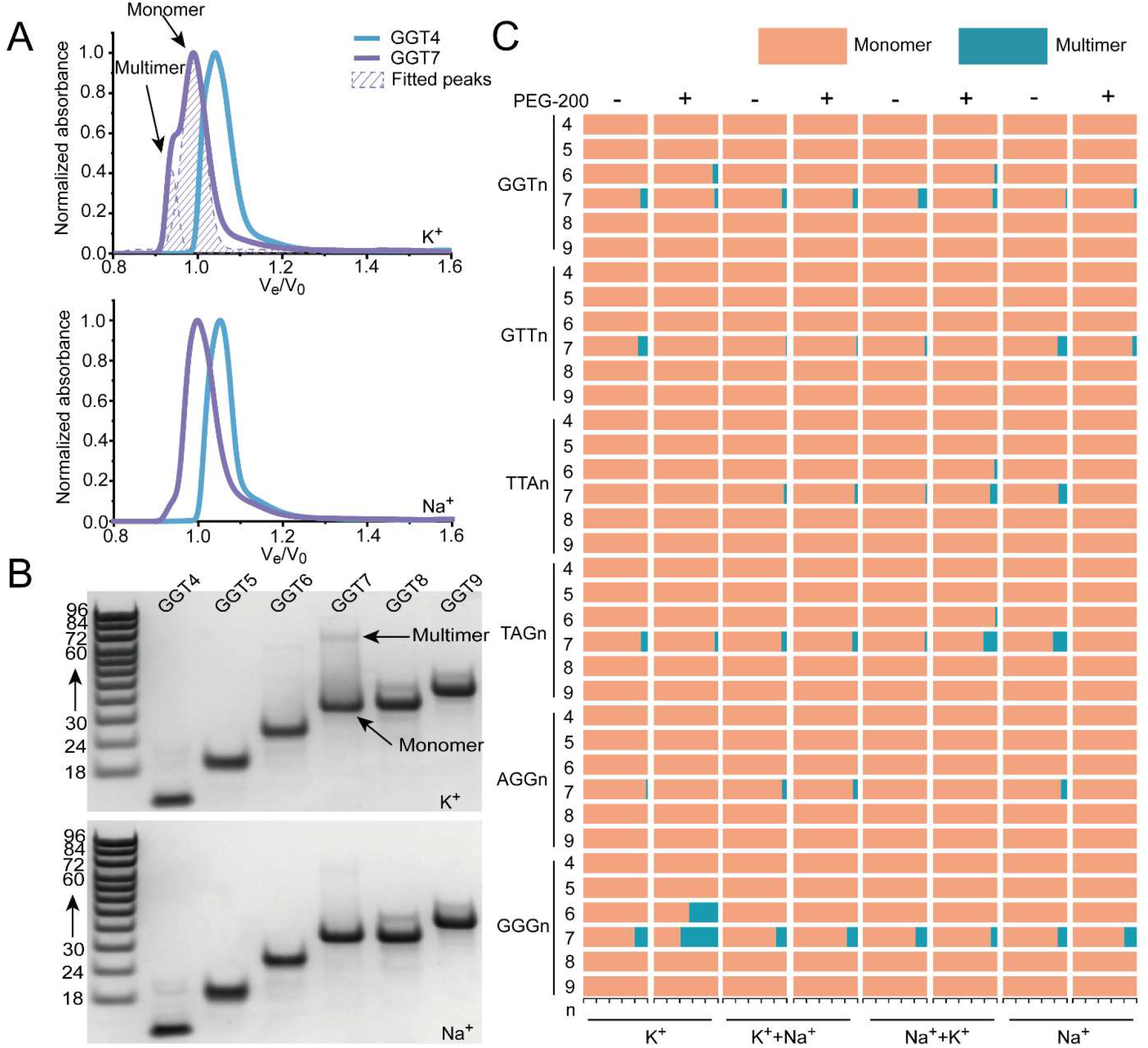
Telomeric G4s Molecularity. (A) SE-HPLC profiles of GGT4 and GGT7 (30 μM) in either K^+^ (up) or Na^+^(down) buffers. (B) Non-denaturing PAGE of GGTn (30 μM) in either K^+^(up) or Na^+^ (down) buffers, using poly(dT_n_) as markers. (C) Summary of proportions of the monomeric and multimer species quantified from the non-denaturing PAGE band.

### Inhibition of DNA polymerase extension by telomeric G4s

To evaluate the replication dynamics under physiologically relevant ionic conditions (100 mM K^+^ and 40 mM Na^+^), DNA polymerase activity was measured by using the six G-overhang permutations with 4 repeats in the template. A 5’-FAM-labeled primer was annealed to templates harboring either the wild-type telomeric G-overhangs (capable of forming stable G4s) or their corresponding mutated sequences (mut-G4, sequences are given in Table S2), in which G to T substitutions were introduced to abolish G4 folding (Figure 5A). Accumulated full-length DNA extension products over time were revealed by denaturing PAGE (Figure 5B for GGT4 and GGG4 and Figure S31 for the four other permutations). Time-course analysis of the extension products via denaturing PAGE revealed a progressive accumulation of full-length DNA strands. Compared to the unstructured mut-G4 controls, phi29 polymerase progression was significantly impeded on the wild-type G4 templates. A statistically significant reduction in full-length product yield emerged on the G4 templates by 10 min (*p <* 0.05) and became highly pronounced at the 30 min endpoint (*p <* 0.001) (Figure 5C, D and Figure S32).

**Figure 5.**
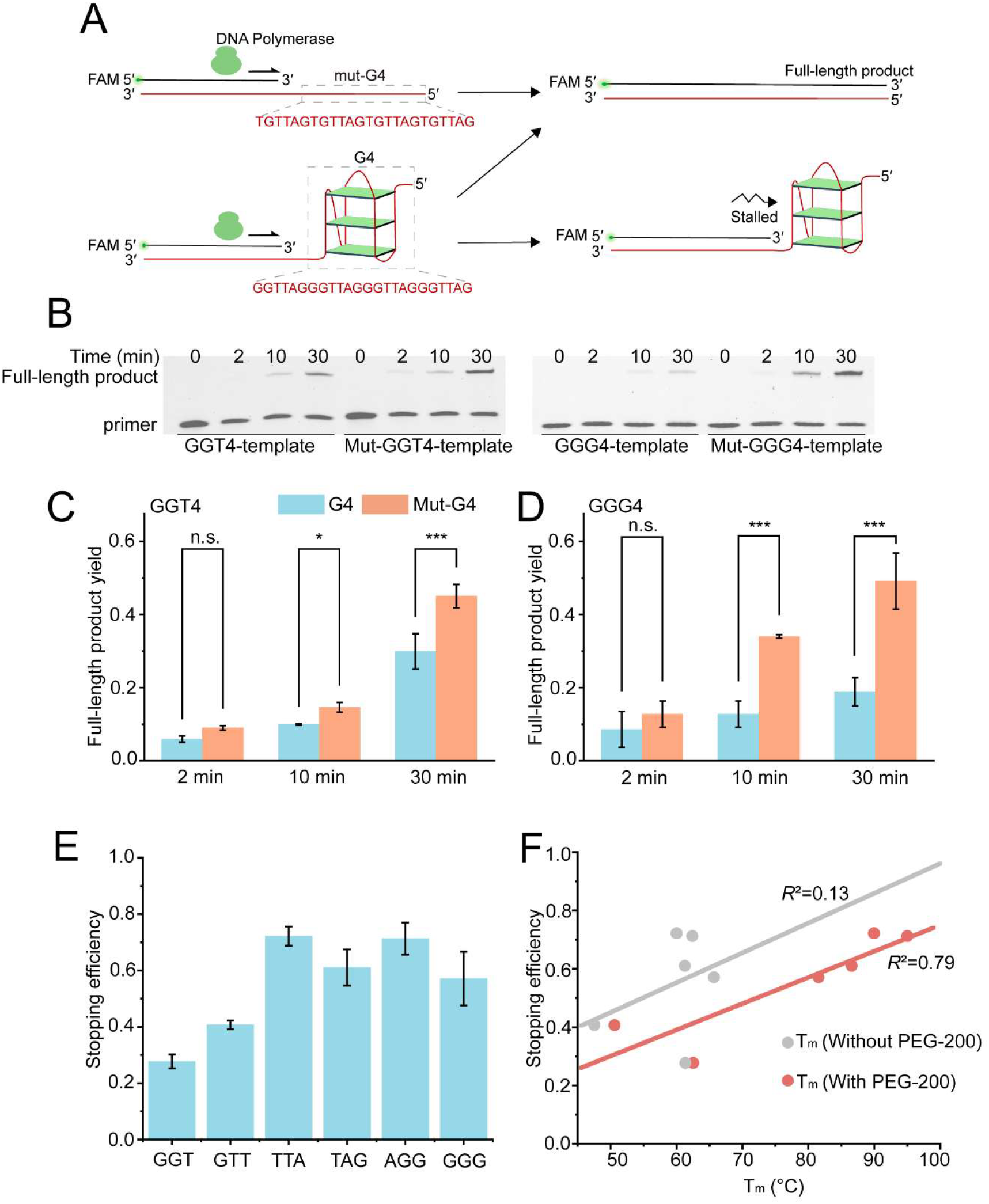
Evaluation of DNA polymerase stalling induced by telomeric G4s. (A) Schematic illustration of the DNA polymerase extension assay on wild-type G4 (G4) and mutated (mut-G4) templates. (B) Denaturing PAGE of the time-dependent extension products catalyzed by phi29 DNA polymerase. Quantitative comparison of full-length product yields for GGT4 (C) and GGG4 (D) at indicated reaction times. The yield defined as [full-length product / (full-length product + unreacted primer)]. (E) Stopping efficiency, defined as 1 – Yield_G4_ / Yield_mut-G4_, at the reaction time of 30 min. (F) Linear regression correlating the stopping efficiency with the T_m_ for G4s formed by six G-overhang in absence and presence of PEG-200. Statistical significance is indicated by * *p <* 0.05, ** *p <* 0.01, and *** *p <* 0.001; n.s., not significant.

To systematically benchmark the inhibitory potency, the stopping efficiency was calculated at a reaction time of 30 min (Figure 5E). G4s formed by all six permutations showed obvious inhibition of DNA polymerase activity (extension efficiency < 1), but exhibited markedly distinct stalling capacities. Consistent with their relatively lower thermodynamic stabilities observed in above UV melting studies, the GGT4 and GTT4 showed a weaker stopping efficiency (∼0.28 and ∼0.41, respectively). Linear regression analysis was performed between the stopping efficiency and the melting temperatures of the respective G4s in the absence or presence of PEG-200. The stopping efficiency showed no correlation with the T_m_ obtained in the absence of PEG-200, but a good correlation in its presence (Figure 5F).

### Rolling C-circle amplification inhibited by the telomeric G4s

A rolling circle amplification (RCA) assay was performed by utilizing a circular DNA template and G-overhang primers with 4 repeats (Figure 6A). The circular template called C-circle was prepared by using the linear sequences of 5′-p-(AACCCT)9-3′ and amplification was checked by non-denaturing PAGE (Figure S33). Mutated G4, named as mut-G4 (in which the second guanine in each G-tract of the wild-type G4 is replaced by thymine to abolish G4 formation), and its corresponding complementary C-circle were used as a control (Figure 6A). The RCA reaction produced ultra-long DNA products, as confirmed by gel analysis (Figure S33). The RCA efficiency was further quantified by a fluorescence enhancement assay [72,73], revealing a pronounced inhibitory effect by five of six telomeric G4s with 4 repeats (Figure 6B). Again, the amplification efficiencies among the six G-overhang permutations (normalized against the mut-G4 yields) exhibit a sequence-dependent behavior (Figure 6B, C). Notably, the GTT4 with the lowest melting temperature exhibited high relative amplification efficiency, further supporting the correlation between reduced G4 stability and attenuated polymerase stalling. While GGG4 also demonstrated a high amplification yield, suggesting that sequence-specific structural dynamics can occasionally override simple thermal stability, the overall trend underscores G4 stability as a primary kinetic barrier. As a comparison, two different protocols (see Materials and methods, Figure 6A) were used to prepare the mixture of C-circle and telomeric G4 before the RCA (Figure 6D, E). Even using protocol P2, five of the six telomeric G4s still showed an obvious inhibition of the RCA efficiency (Figure 6D). Interestingly, varying the thermal preparation from 37 °C incubation (P1) to 95 °C annealing (P2) resulted in highly sequence-dependent responses (Figure 6E). While GGG4 and AGG4 exhibited comparable amplification efficiencies with minimal disparity, the other permutations displayed significant differences. This distinct sequence-dependent thermal response suggests that for certain permutations (such as GTT4), high-temperature annealing may actually favor the folding of robust G4s rather than the amplification-favorable primer-template duplex, thereby further inhibiting polymerase extension. Overall, these results indicate that telomeric G4s can act as potent barriers to continuous polymerase progression, whether serving as primers (Figure 5) or as templates (Figure 6).

**Figure 6.**
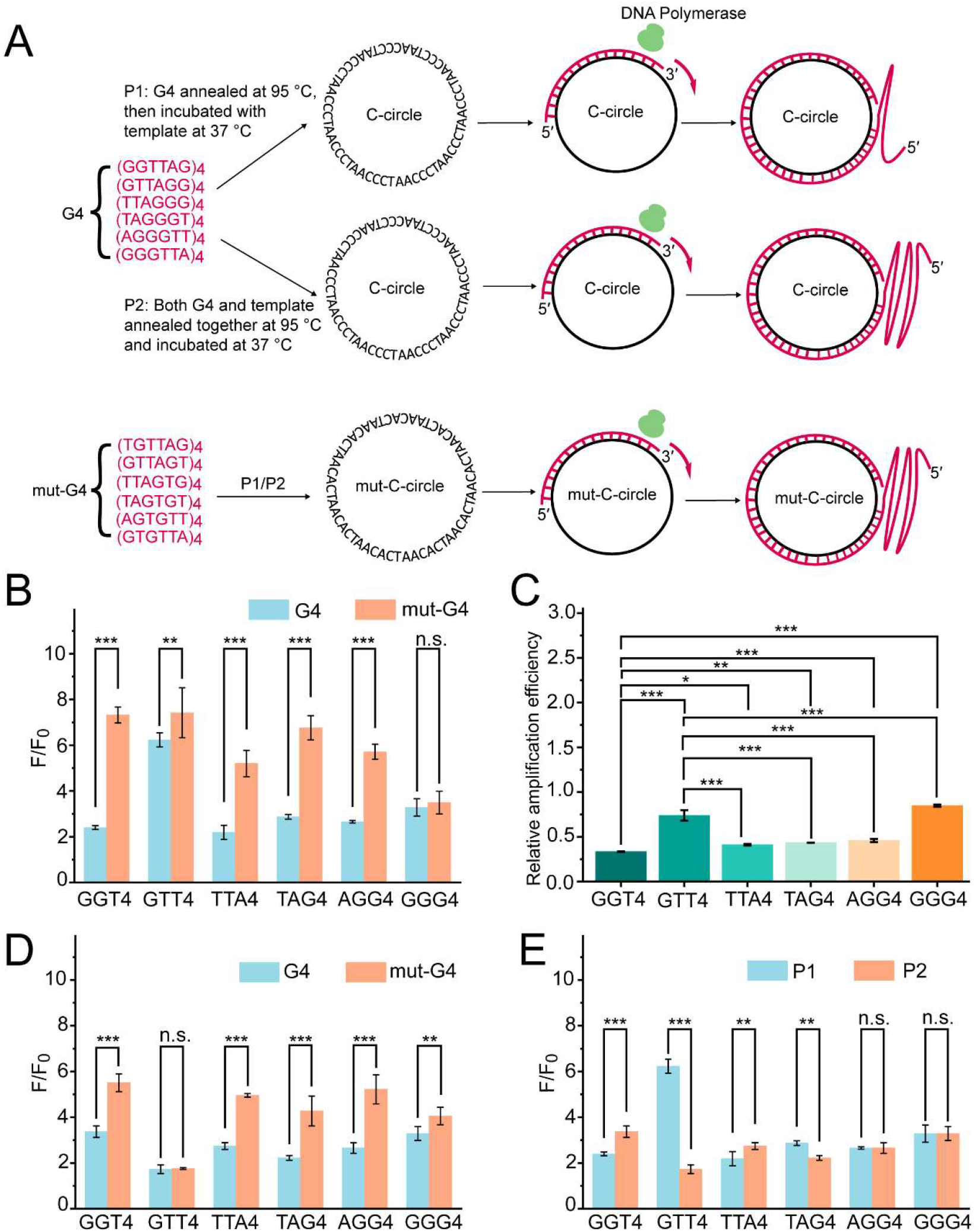
Impact of telomeric G4s on RCA efficiency. (A) Schematic illustration of the RCA assay utilizing a circular DNA template (C-circle) primed by a wild-type G4 or mutated (mut-G4) sequence. (B) Quantitative comparison of the RCA yields. The F and F_0_ represent the fluorescence intensities of SYBR-Gold with and without the RCA products, respectively. (C) Relative amplification efficiencies, defined as [(F/F_0(G4)_) / (F/F_0(mut-G4)_)], among the six G4 permutations using protocol P1. (D) Quantitative comparison of the RCA yields using protocol P2. (E) Quantitative comparison of the RCA yields for the wild-type G4 using protocols P1 and P2. Statistical significance is indicated by * *p <* 0.05, ** *p <* 0.01, and *** *p <* 0.001; n.s., not significant.

## Discussion

Recent nucleotide-resolution measurements reveal that human telomeric G-overhangs terminate predominantly in 5′-GGTTAG-3′ (61%) and 5′-GTTAGG-3′ (13%) [54]. This non-random distribution is intrinsically governed by telomerase catalysis and chromosome end-processing mechanisms. During telomere extension, human telomerase regulates processivity via a sequence-defined pause signal embedded within the hTR template [74]. Following the addition of the sixth nucleotide (dG6), the resultant dT:rA base pair creates a kinetic barrier characterized by a significantly elevated K_m_ for the subsequent dG1 incorporation. This slow dG1 addition step promotes complete product release exactly at this physical boundary, thereby yielding nascent DNA generally terminating in the GGTTAG register [75]. Biologically, this predominant GGTTAG terminus functions in evolutionary concert with the shelterin complex. Crystal structures reveal that the single-stranded DNA-binding protein POT1 specifically recognizes the decameric sequence TTAGGGTTAG, tightly sequestering the 3′-terminal guanines within a basic binding pocket to avert aberrant DNA damage signaling and nucleolytic degradation [76]. Consequently, overhang permutations generated by incomplete processing may undergo continuous enzymatic resection until this optimal GGTTAG boundary is exposed and stably protected.

Despite these well-characterized biosynthetic origins, the intrinsic structural behaviors of the diverse hexanucleotide permutations have been largely overlooked, as prior G4 studies have disproportionately focused on the canonical contiguous (TTAGGG)_n_ tract [59,60]. Our comprehensive biophysical analysis demonstrates that all six naturally occurring terminal permutations, including the highly abundant GGTTAG and GTTAGG permutations harboring a split G-tract, spontaneously fold into stable G4 structures (Figures S1-7). Crucially, the subtle positional variations of the nucleotides within these distinct registers dictate markedly different G4 topologies (Figure 2) and thermodynamic stabilities under both potassium and sodium environments (Figure 3). This acute sensitivity to the cationic balance aligns with a recently described structural switch mechanism, where minor sequence alterations dictate distinct folding equilibria in response to specific alkali metal ions [77]. Consequently, our findings reveal a previously unrecognized dimension of sequence-dependent structural regulation at chromosome ends.

Firstly, telomeric permutations containing split G-tracts (GGTn and GTTn) are capable of folding into stable G4s. While G4s formed by all contiguous G-tract sequences generally exhibit robust thermal stability, the two split G-tract permutations display divergent thermodynamic profiles. The GTTn series exhibits significantly lower melting temperatures, whereas GGTn remains unexpectedly stable (Figure 3). The severe destabilization of GTTn likely arises either from the loss of a G-quartet or, assuming 3 quartets are still formed, from the energetic penalty incurred when guanines must be recruited from distant adjacent repeats to complete the G-tetrad core, which forces the formation of extended and energetically unfavorable loop architectures. The disruption of the continuous guanine register fundamentally alters the global folding topology. During the preparation of this manuscript, a nuclear magnetic resonance structural elucidation demonstrates that splitting the initial G-tract of GTT4 forces the telomeric G4 to adopt a two-layer antiparallel topology with K^+^ (PDB: 9I5S) [78]. Consistent with this unique structural constraint, our circular dichroism data reveal that GTT4 uniquely adopts and firmly retains an antiparallel topology across all tested ionic and crowding conditions (Figure 2). It strongly resists the typical hybrid to parallel structural transitions that are universally observed in contiguous repeats under dehydrating environments [78,79]. Consequently, the precise position of the split G-tract dictates a delicate balance between thermodynamic stability and conformational plasticity for these naturally abundant overhang permutations.

The intracellular environment is highly crowded, which fundamentally alters the thermodynamics of nucleic acid folding compared to dilute solutions. Consistent with previous biophysical investigations, the introduction of PEG-200, proposed to mimic crowding, globally enhanced the thermal stability of the telomeric G4s (Figure 3B) and promoted a topological shift toward parallel conformations (Figures 2 and S17) [23,80]. However, this crowding induced structural transition is predominantly driven by local dehydration effects rather than pure steric exclusion [81]. Intriguingly, for permutation repeat numbers of 7 and 6, the addition of PEG-200 induced an obvious biphasic melting and annealing behavior (Figure 3). Unlike the monophasic melting universally observed in dilute buffers, this stepwise thermal transition indicates an increased conformational heterogeneity under crowding conditions. A similar phenomenon was observed with G4s formed by long Oxytricha telomeric DNA repeats (TTTGGGG)_n_ [23]. Concentration-dependent melting experiments showed that the lower-melting temperature arises from multimeric species, whereas the higher-melting temperature originates from intramolecular species (Figure 4) [23].

Human telomeric G-overhang spans hundreds of single-stranded nucleotides and typically fold into higher-order G4s resembling a beads-on-a-string architecture [82]. This higher-order G4 structure formed to promote maximal G-quartet formation and relied on G4s containing entirely contiguous G-tracts. While both GGT8 and GTT8 contain a split G-tract, GGT8 but not GTT8 exhibit a similar beads-on-a-string architecture. The melting temperature of GGT8 in K^+^ buffer is 9.2 °C lower than that of GGT4, which closely matches the thermal stability difference seen between GGG8 and GGG4 (Figure 3 and Table S3). This similarity implies that the G4s formed by GGT8 and GGT4 may share an equivalent number of stacked G-quartets. In addition, both GGT4 and GGT8, or GGG4 and GGG8 fold into an intramolecular G4 with a hybrid conformation in K+ buffer (Figures 2 and 4), indicating that GGT8 may fold into two non-interacting hybrid G4s with a TTA linker. In contrast, GTTn behaves differently: GTT4 forms antiparallel intramolecular G4s, yet GTT8 adopts a hybrid topology in K^+^ buffer (Figures 2 and 4). The comparable melting temperatures of GTT4 and GTT8 (Figure 3 and Table S3) further suggest that the additional guanines either contribute to G-quartet formation or that inter-G4 interactions compensate for the potential energetic penalty [83]. Such architectural divergence at the higher-order level offers a potential target for pharmacological intervention [84]. Specifically, the distinct interfaces formed between adjacent G4s and the unique loop conformations dictated by split G-tracts could yield highly specific structural pockets suitable for ligand binding.

The consequence of the observed thermodynamic or structural disparities is demonstrated by DNA polymerase extension assays (Figure 5A) [85]. The folded G4 structures serve as potent kinetic barriers to DNA polymerase progression (Figure 5B, C, D). The stopping efficiency was strongly dependent on the nature of the sequence (Figure 5E). Consequently, permutations containing split G-tracts exhibit significantly attenuated stalling capacities compared to the highly stable contiguous G-tract permutations. Furthermore, a linear correlation between stopping efficiency and melting temperatures is observed when thermodynamic stabilities are evaluated under molecular crowding conditions, but not in dilute conditions (Figure 5F). These findings indicate that the intrinsic destabilization of the naturally abundant split G-tract termini may lower the kinetic barrier for replication machineries. Beyond the intrinsic kinetic barriers encountered by polymerases, the complete resolution of telomeric G4s in vivo requires specialized helicases to prevent fork stalling and genomic instability [86]. The distinct topological features and generally lower stabilities of the dominant split G-tract permutations (*e*.*g*., GGTTAG) suggest that these naturally abundant sequences may serve as thermodynamically favorable substrates for these helicases, thereby optimizing local DNA unwinding kinetics during replication.

In addition to regulating replication, G4 folding creates a native steric barrier that modulates access of telomerase and other telomeric proteins [34]. The dominant split G-tract permutations form less stable G4s than their contiguous repeats. Consequently, their structural equilibrium is likely shifted toward a dynamic unfolded state. This shift may facilitate the loading of single-stranded binding proteins such as POT1, which sequence-specifically sequesters the unfolded DNA to recruit and regulate telomerase [87]. Furthermore, the specific G4 folding preferences of these permutations under molecular crowding may also govern the formation of telomeric DNA: RNA hybrids (R-loops), a critical structural driver of homologous recombination in ALT-positive tumors [52,53].

In conclusion, our systematic analysis demonstrates that characterizing human telomeres exclusively through the canonical TTAGGG repeat oversimplifies their structural landscape. The G4 structures formed by split G-tracts within the most abundant chromosomal termini (GGTTAG and GTTAGG repeats) exhibit distinct stabilities, topologies, higher-order assembly behaviors, and polymerase stop efficiencies. Recognizing this expanded scope of sequence diversity provides critical insights into telomere regulation. Ultimately, potential therapeutic strategies, whether aimed at stabilizing G4, displacing POT1, inhibiting telomerase, or increasing replication stress in ALT cancers, should consider the unique structural features of these dominant split G-tract permutations to enable precise targeting and optimal clinical efficacy.

## Supporting Information

The Supporting Information is available free of charge. Contents include: lists of currently resolved human telomeric G-quadruplex structures and oligonucleotides used in this study; summary of T_m_ values and comprehensive UV melting/annealing curves; thermal difference spectra (TDS) and circular dichroism (CD) spectra detailing conformational changes and r values under various ionic and crowding conditions; fluorescence enhancement profiles using ThT and NMM probes; size-exclusion chromatography (SE-HPLC) and native PAGE analyses assessing G4 molecularity and multimerization; and supplementary gel electrophoresis and quantitative data for the DNA polymerase extension and rolling circle amplification (RCA) assays.

## Acknowledgements

This work was supported by the National Natural Science Foundation of China (22307143), National Natural Science Foundation of China for Excellent Young Scientists Fund Program (Overseas), National Foreign Experts Program (S20250128), Natural Science Foundation of Jiangsu Province, China (BKA20231007), Innovation and Entrepreneurship (Shuangchuang) Program of Jiangsu Province (JSSCTD202447) and Jiangsu Province Frontier Technology Research and Development Program (BF2024075). J.L.M. acknowledges recurrent funding from CNRS, Inserm and Ecole Polytechnique.

